# DNA polymerase stalling at structured DNA constrains the expansion of Short Tandem Repeats

**DOI:** 10.1101/2020.06.20.162743

**Authors:** Pierre Murat, Guillaume Guilbaud, Julian E. Sale

## Abstract

**Background:** Short tandem repeats (STRs) contribute significantly to *de novo* mutagenesis, driving phenotypic diversity and genetic disease. Although highly diverse, their repetitive sequences induce DNA polymerase slippage and stalling, leading to length and sequence variation. However, current studies of DNA synthesis through STRs are restricted to a handful of selected sequences, limiting our broader understanding of their evolutionary behaviour and hampering the characterisation of the determinants of their abundance and stability in eukaryotic genomes.

**Results:** We perform a comprehensive analysis of DNA synthesis at all STR permutations and interrogate the impact of STR sequence and secondary structure on their genomic representation and mutability. To do so, we developed a high-throughput primer extension assay that allows monitoring of the kinetics and fidelity of DNA synthesis through 20,000 sequences comprising all STR permutations in different lengths. By combining these measurements with population-scale genomic data, we show that the response of a model replicative DNA polymerase to variously structured DNA is sufficient to predict the complex genomic behaviour of STRs, including abundance and mutational constraints. We demonstrate that DNA polymerase stalling at DNA structures induces error-prone DNA synthesis, which constrains STR expansion.

**Conclusions:** Our data support a model in which STR length in eukaryotic genomes results from a balance between expansion due to polymerase slippage at repeated DNA sequences and point mutations caused by error-prone DNA synthesis at DNA structures.

## Background

Nearly half of the human genome is composed of various forms of DNA repeat, which contribute to gene function, genome structure and evolution [1]. Among DNA repeats, motifs of 1 to 6 nucleotides repeated in a head-to-tail manner, known as Short Tandem Repeats (STRs) or microsatellites, have attracted extensive attention due to their highly polymorphic nature and consequent applications in forensic DNA fingerprinting, genetic linkage analysis and the study of population dynamics [2]. STRs are ubiquitous in eukaryotic genomes, with ∼4,500,000 loci covering up to 2.5% of the human genome [3]. They exhibit mutation rates that are orders of magnitude higher than for other variant types such as single nucleotide polymorphisms (SNP) or copy number variations [4, 5]. STR length variation in genes, particularly expansion, is linked to the aetiology of various human neurodegenerative diseases [6]. Recently, length polymorphism of STRs has been associated with epigenetic plasticity [7] and gene expression variation [8, 9]. However, the mechanisms driving and constraining the evolution of STRs length are poorly understood.

STR length variation is generally thought to arise from replication slippage events during which the nascent DNA dissociates from its template and then reanneals out of register, an event facilitated by the repetitive nature of STRs [2]. The resulting insertion or deletion of repeat units is countered by repair pathways, notably mismatch repair, but a small fraction of events become fixed and transmitted across cell division. The low complexity nature of STRs also makes them prone to fold into intrastrand, non-B form secondary structures that may impede DNA polymerase progression. Inaccurate resolution of the resulting replication intermediates may also lead to length variation [10]. While mismatch repair (MMR), base/nucleotide excision (BER and NER) and post-replication repair (PRR) proteins can all modulate STR length variation in a wide variety of model systems [11], it has been demonstrated *in vitro* that DNA polymerase alone is able to generate STR expansions through slippage [12].

It is generally believed that the propensity of STRs to form secondary structures drives their genomic instability [13, 14]. This assumption largely stems from the study of large trinucleotide repeat expansions involved in disorders such as Huntington’s disease or Fragile X syndrome [6]. *In vitro* biophysical characterisation of these repeats revealed a correlation between their ability to form hairpin-like structures, and the observed sequence and length dependence of expansion [15–17]. These observations prompted the structural characterisation of STRs involved in other diseases and identified several STR motifs prone to fold into non-B form secondary structures. For example, the tetranucleotide (CCTG•CAGG) repeat, which contributes to the genetic instability associated with Myotonic Dystrophy Type 2, also folds into an hairpin-like structure [18], the hexanucleotide repeat (GGGGCC), expansion of which triggers familial amyotrophic lateral sclerosis (ALS) and frontotemporal dementia (FTD), adopts a G-quadruplex (G4) structure [19] and the hexanucleotide (CCCCCG) repeat found within the promoter of the human DAP gene forms i-motifs under physiological conditions [20]. Nevertheless, the apparent enrichment of non-B form secondary structures within long disease-related STRs could reflect that the structures have a deleterious impact on repair mechanisms [11], rather than them being a causative factor for expansion. Moreover, the proposed correlation between structure and length instability is based on the study of selected and pathologic STRs with no evidence that the conclusions are generalisable across all STR motifs.

Several mutation models have been proposed to explain the equilibrium distribution of STR lengths [21]. Traditional models, such as the Stepwise Mutation Model [22], have been proven useful to analyse differences in STR length between individuals. However, it fails to explain why individual STR loci do not expand indefinitely. An alternative model proposed that repeat lengths at equilibrium result from a balance between slippage events and point mutations [23]. This model relies on a balance between the rates of slippage and point mutations that limit the length of perfect repeats. Together with the observation that the rate of contractions increases exponentially with repeat length in humans [21], these models predict a maximum length for perfect STRs at equilibrium. Nevertheless, these and even more advanced models, which are able to quantify the magnitude of the length constraint for a given STR motif [24], fail to explain the origin of these selective constraints or the factors that determine the abundance and behaviour of the different STR motifs in eukaryotic genomes [25].

In this study, we set out to address the impact of DNA structures on DNA synthesis at STRs in order to understand the relationship between the ability of a given STR to stall DNA polymerase and its genomic stability. To do this, we developed a high-throughput polymerase extension assay that allowed us to monitor the kinetics of DNA synthesis at all STR permutations in different lengths, in parallel. We have used the assay to map at single-nucleotide resolution the movement of a prototypical replicative DNA polymerase through the repeats over time. From this kinetic data we are able to infer the secondary structure adopted by a given STR and link this to slippage and point mutation during DNA synthesis. We selected the A-family T7 DNA polymerase (Sequenase) as a model polymerase in our assay for its high processivity and lack of exonuclease activity in order to detect within the course of a single round of DNA synthesis all possible deleterious effect of DNA structures on DNA synthesis efficiency and fidelity without the need for additional accessory factors. We demonstrate that the response of DNA polymerase to variously structured DNA is sufficient to predict the complex genomic behaviour of STRs, including abundance and mutational constraints. We show that structured STRs exhibit lower relative abundance and shorter length, suggesting they are generally deleterious for eukaryotic genomes. Unexpectedly, the lengths of structured STRs are also globally more stable over time. This greater length constraint is imposed by more frequent point mutations limiting the extent of the perfect repeats. We propose a model in which the distribution of STR lengths at equilibrium results from the ability of DNA structures to induce error-prone DNA synthesis that limit STR expansion. These observations have important implications for understanding the evolution of STRs and for appreciating how they shape eukaryotic genomes.

## Results

### Pooled measurement of DNA polymerase stalling at designed structured DNA sequences

To address how DNA secondary structures impede DNA synthesis, we designed a library of 20,000 sequences (Additional file 2) and devised a method for accurately measuring polymerase stalling at these sequences in a single experiment. The library comprises all 5,356 possible STR permutations of 1-6 nucleotides in three different lengths (24, 48 and 72 nt, giving a total of 16,068 sequences). The library also contains positive control sequences, designed to fold into known single-stranded secondary structures such as hairpins (960 sequences), G4s (1500 sequences) and i-motifs (472 sequences), of various lengths and GC contents in order to cover potential DNA structures of a wide range of thermodynamic stabilities (see Methods). In order to control that polymerase stalling is due to the structure rather than the GC content of a sequence, we also included negative control sequences in the form of 1,000 random sequences of varying GC content (from 20 to 80%) (see Methods details). We obtained the library as a mixed barcoded oligonucleotide pool synthesized on, and eluted from, a programmable microarray and inserted it into a phagemid vector (Fig. 1a). The library was then amplified in *E*. *coli* and circular single-stranded DNA templates containing the library were produced using M13KO7 helper phage.

**Fig. 1.**
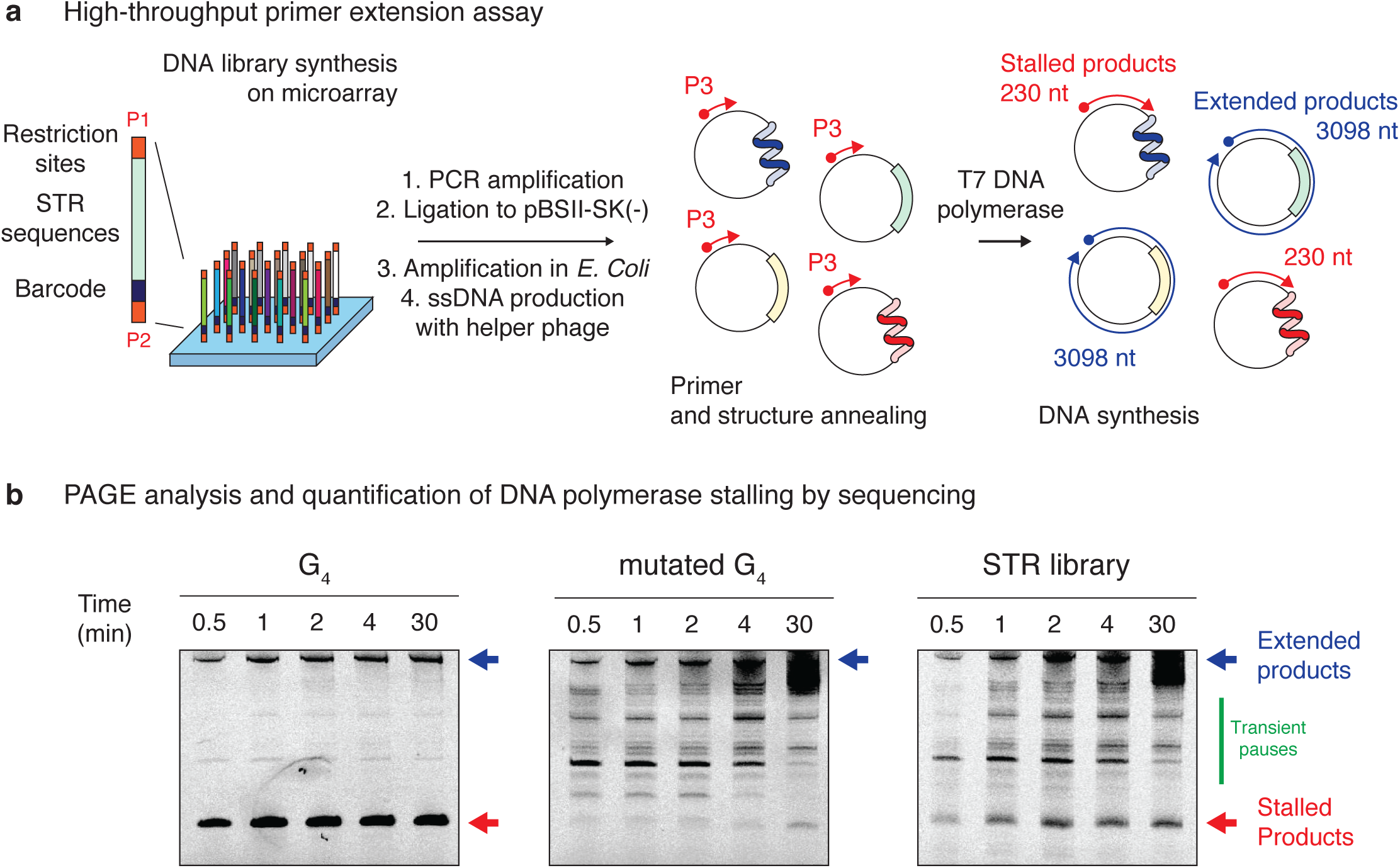
Pooled measurement of DNA polymerase stalling at STRs. **(a)** Overview of the high-throughput primer extension assay used to monitor DNA synthesis at designed sequences. A library of 20,000 sequences comprising all STR permutations at three different lengths together with control structured DNA sequences was synthesised on a programmable microarray, eluted and inserted into a phagemid vector. After PCR amplification, insertion into a phagemid vector and bacterial amplification, circular single-stranded DNA templates were produced using a M13KO7 helper phage. Fluorescently labelled primer (P3) and structures annealing were performed before initiating DNA synthesis through the addition of T7 DNA polymerase. Primers are then either fully extended to the length of the circular template or the extension is stopped within STRs if the DNA polymerases stall at structured DNAs. **(b)** Extended and stalled products were then analysed by denaturing Poly Acrylamide Gel (PAGE) electrophoresis, recovered from the gel matrix and prepared for high throughput sequencing. DNA polymerase stalling was then quantified by analysing the enrichment of each sequence form the library in the stalled and extended fractions. Representative fluorescence gel imaging of primer extension reactions on templates containing a G-quadruplex (G4) structure, a mutated G4 or the entire DNA library, stopped after the indicated times, are reported for comparison. Blue and red arrows indicate the position of the extended and stalled products respectively. The green line highlights the presence of transient stall sites that disappear overtime.

We performed a primer extension assay on these templates using a fluorescently labelled primer annealed ∼200 nt from the 3’ end of the designed sequences [26]. The primer was annealed under conditions favouring the formation of secondary structures within the single stranded templates (see Methods details). The reactions was initiated by the addition of a modified T7 DNA polymerase [27] (Sequenase), which was selected as a model A-family replicative polymerase [28] that exhibits high processivity and high fidelity in the absence of proofreading/exonuclease activity.

The stalled (∼ 200 nt) and extended (∼ 3000 nt) products at different time points (0.5, 1, 2, 4 and 30 min) were separated by electrophoresis, their positions having been first determined by using templates containing a G4 motif or mutated control not expected to support G4 formation (Fig. 1b). With the G4 template, stalled products accumulated at the position of the G4 motif and persisted over the time course of the experiment. At the end of the primer extension assay, the intensity corresponding to the stalled product represented 70 % of the total signal indicating that the G4 structure significantly impedes polymerase progression (Fig. 1b left panel). Conversely, only transient replication pauses at other sites in the template (but not within the inserted library), which were resolved within the course of the experiment, were observed when using the mutated G4 template. The fully extended and stalled products for each time point were recovered from the gel matrices and prepared for high-throughput sequencing (see Methods).

In order to quantify the ability of a given sequence to stall DNA polymerase, we computed a stall score, hereafter refer to as σ, as the ratio of the normalised number of reads assigned to this sequence in the stalled products over the total normalised number of reads in both the stalled and extended fractions. Hence the stall score is a value between 0 and 1 with higher values reflecting a greater ability of a given sequence to stall polymerase progression, a value of 1 indicating the presence of a given sequence in the stalled fraction only. We found that the pipeline itself does not introduce any sequence representation bias and allows us to compute reproducible stall scores for each sequence from the DNA library (Spearman Correlation = 0.722 between replicates, Additional file 1: Figure S1).

### The kinetics of DNA synthesis highlights structure-dependent transient and persistent stalling events

We first examined the performance of our method for monitoring the kinetics of DNA synthesis through known secondary structure-forming sequences. We computed the distribution of the stall scores for the control sequences and of the STR library at each time point (Fig. 2a-c, Additional file 1: Figure S2a and S2b). While the stall scores of designed structured sequences were globally higher than those from the negative control sequences (random sequences of varying GC content) at each time point (*P* < 1.4 x 10^−7^, Additional file 1: Figure S2c), the scores are highly structure and time-dependent. For example, stall scores of G4-forming sequences displayed a median at 0.48 at 0.5 min and 0.83 at 30 min (Fig. 2a). In contrast, the stall scores of hairpin-forming sequences showed a different trend with scores shifting to lower values over time from 0.40 at 0.5 min to 0.13 at 30 min (Fig. 2b). The stall scores associated with the STR sequences display a more complex pattern, which suggest that STRs fold into a wide range of structures. It is noteworthy that a significant portion of the STR sequences displayed stall scores of 1, *i*.*e*. they are only in the stalled and not extended fraction (Fig. 2c). It is important to remember that the stall score of a given sequence at a given time point is determined relative to all the other sequences in the library, *i*.*e*. stall scores are relative rather than absolute values. Hence if the stall score associated with a sequence varies over time, all other stall scores will also change as the normalised total number of reads remains constant. Thus, this suggests that while both types of sequence stall DNA synthesis at 0.5 min, the stall resulting from G4s is more likely to be persistent, while those from hairpin-forming sequences are more likely to be resolved. Additionally, we found that stall score correlated well with the predicted stability of the designed structures (Additional file 1: Figure S2d-f). Taken together, these observations demonstrate that stalling at the sequences designed to fold into secondary structures is due to their structure rather than their GC content and that the ability of the polymerase to resolve the stall depends on the nature of the DNA structure.

**Fig. 2.**
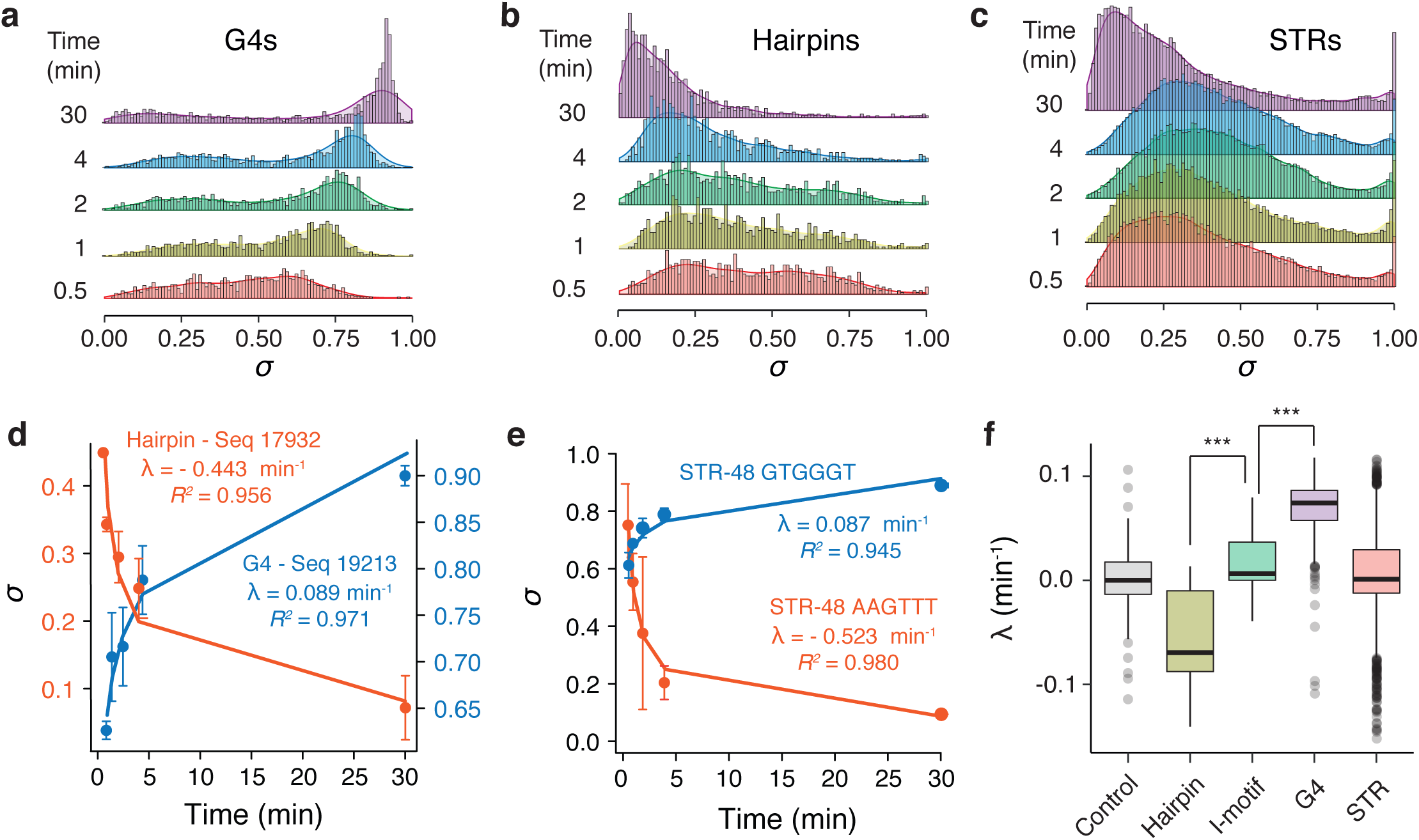
Monitoring the kinetics of DNA synthesis reveals structure-dependent transient and persistent stalling events. Distribution and density plots of the computed stall scores associated to the **(a)** G4-, **(b)** hairpin-forming control sequences and **(c)** all STR permutations at different time points showing time-dependent DNA polymerase stalling at DNA structures. **(d)** Example of time-dependent variation of stall scores associated to a G4 (<u>GGGG</u>ACA<u>GGGGGG</u>AC<u>GGGG</u>CTTGAAAT<u>GGGG,</u> blue points) or a hairpin (<u>TACGGTATAATTAAGGACGTAT</u>TTTT<u>ACGTCCTTAATTATACCGTA,</u> red points) structure. The kinetic constants associated with DNA synthesis (*λ*) together with their associated coefficients of determination (*R*^*2*^) were extracted by fitting the experimental data using either an exponential growth or decay functions. Error bars represent the standard deviation from two independent replicates. **(e)** Similar analysis performed on two representative STRs, the GTGGGT and AAGTTT sequences repeated 8 times, highlighting transient or persistent stalling at STRs. **(f)** Distribution of the kinetic constants of DNA synthesis associated with the control sequences, *e*.*g*. random sequences of varying GC content (control), hairpins, i-motifs and G4s; and all STR permutations. λ values associated with the hairpins, i-motifs and G4s are for structured sequences that have a stall score statistically higher than the random control sequences at each time point. Centre lines denote medians, boxes span the interquartile range, and whiskers extend beyond the box limits by 1.5 times the interquartile range. *P* values for the comparison of the distributions were calculated using Kolmogorov–Smirnov tests, \*\*\**P* < 0.001.

We reason than following the evolution of sigma over 5 time points would results in a robust measurement as the change in stall score over time, which reflects the kinetics of stalled polymerases resolution, followed either exponential growth or decay functions, consistent with the structures impeding DNA synthesis by triggering dissociation of the DNA polymerase from its template [29]. We thus extracted the kinetic constants; hereafter refer to as λ, associated with each sequence present in the DNA library using exponential growth/decay models. While G4 of high stall scores are characterised by positive λ values, *i*.*e*. exponential growth functions with respect to time, hairpins are characterised by negative λ values, *i*.*e*. exponential decay functions (see Fig. 2d for examples). These observations suggest that the response of the polymerase to these structures is quite different: stalling at a hairpin is transient, stalling at a G4 is persistent. As expected by the diversity of potential forming structures, we found both negative and positive constants for the STR sequences, (Fig. 2e). We then analysed the distribution of λ values associated with each of our rationally designed control structured sequences that have a stall score significantly higher than the random control sequences at each time point. While for 89.4% of sequences forming hairpins λ was negative, 97.6% of those forming tetrahelical structures, *i*.*e*. G4s and i-motifs, had a positive λ, confirming the transient or persistent stalling at hairpin and tetrahelical DNA structures respectively. Importantly, each structural class globally exhibits λ constants that are statistically different to each other (Fig. 2f). This observation suggests that λ, which reflects the evolution of the stall score, σ, over time can be used to distinguish between these DNA structures. It is worth noting that the λ constants correlate with the predicted stabilities of the structures (Additional file 1: Figure S2g-i) showing that structure formation, rather than the DNA sequence, affects the kinetic of stall resolution. Nevertheless, because the λ constants associated with the hairpin and tetrahelical structures span the range of the values associated with the negative controls, additional information is needed to infer STR structures. The λ values associated with STRs are centred around 0 but span the range of values observed for hairpins, G4s and i-motifs (Fig. 2f) suggesting that the STR library contains sequences capable of forming both hairpin-like and tetrahelical structures.

### Inferring STR structures from polymerase stalling events

Because the intrinsic secondary structure of an STR may dictate its behaviour in eukaryotic genomes, we aimed to assign a structural class to each STR motif based on polymerase response. To do this, we devised a supervised machine-learning pipeline to predict the structural class of all STR motifs from the experimental data generated by our polymerase extension assay (see Methods details). We used the control sequences to train a classifier algorithm to assign one of the 4 following classes: hairpins, G4s, i-motifs and unfolded (referred to as HAIRP, QUAD, IMOT and UNF hereafter) to each of the 964 unique single-stranded motifs from our primer extension assay measurements. Sequences falling into the UNF class are sequences whose stall scores at each time point were not statistically different from the stall scores of the random negative control sequences, *i*.*e*. sequences that are not structured under the condition of our polymerase extension assay. It is noteworthy that the most important features for assigning a structure to each STR motif were the *P* values associated with the statistical differences between the stall score (σ) of the STR and the control sequences and the constant, λ (Additional file 1: Figure S3b and S3c), indicating that the strength of polymerase stalling and the kinetics of DNA synthesis are the most informative features for predicting the structure of the STRs. Fig. 3a reports the stall scores of each STR single-stranded motif together with their assigned structural classes when 48-nt long (results for other motif lengths are reported in Additional file 1: Figure S4a and S4b). These figures allow identification of STR motifs with high stall scores and their predicted structures falling into the UNF, HAIRP, IMOT and QUAD structural classes.

**Fig. 3.**
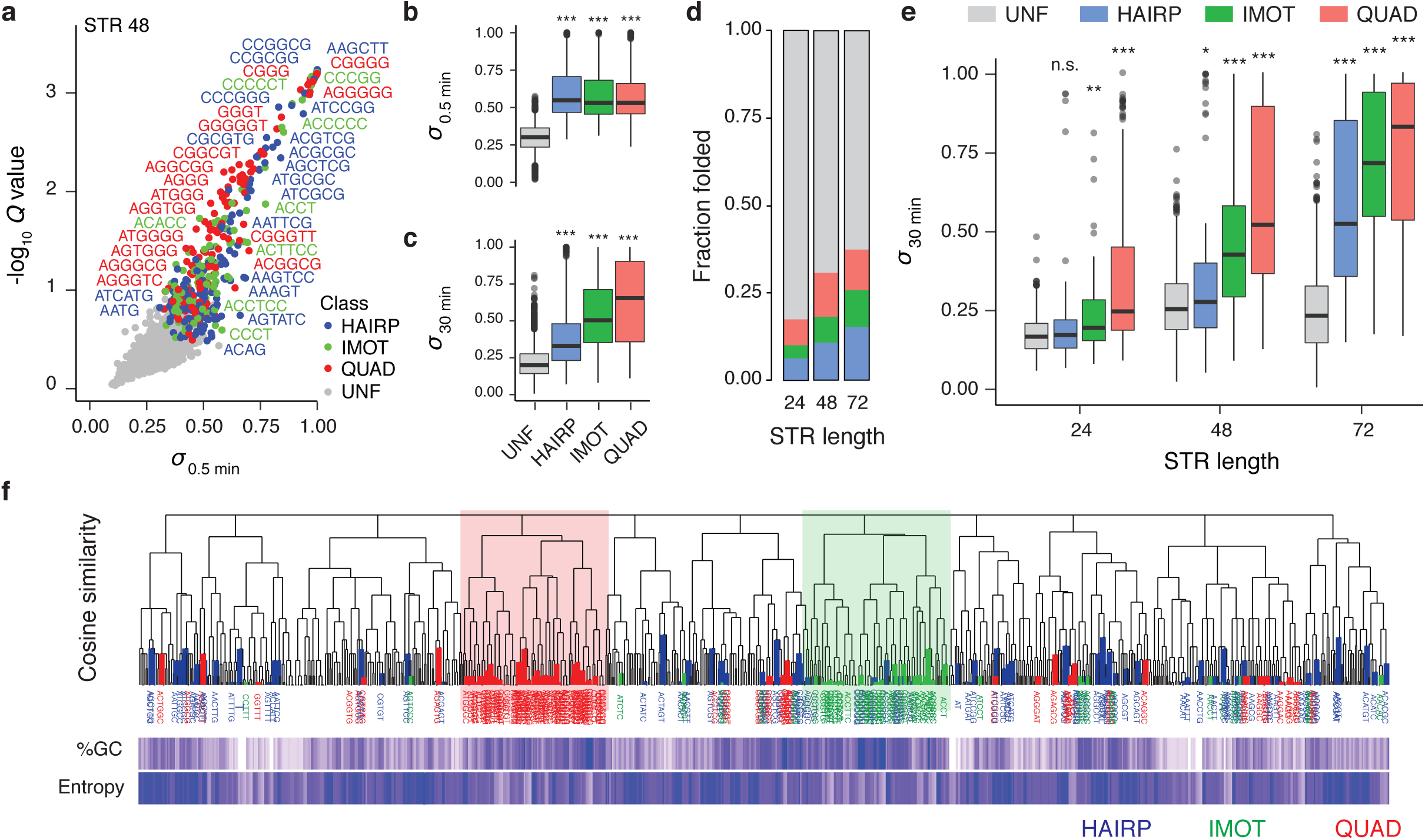
Hairpin-like and tetrahelical structures arise from distinct STR family with distinct properties. **(a)** Distribution together with the statistical relevance of averaged computed stall scores at 0.5 min for the 964 unique single-stranded STR motifs when 48 nucleotides long together with their structures predicted using a supervised machine-learning algorithm. The *Q* values are computed by combining *P* values associated to the statistical differences in stall scores at all time points observed for a given STR motif and the random control sequences of varying GC content (see Methods details) and reflect that DNA polymerase stalling at STRs is structure rather than sequence dependent. Distribution of stall scores at **(b)** 0.5 min and **(c)** 30 min associated with the different structural classes of STRs of any length, highlighting the transient or persistent nature of DNA polymerase stalling at hairpin-like and tetrahelical STR respectively. **(d)** Fraction of predicted structured STRs as a function of STR length highlighting length-dependent structure formation. **(e)** Distribution of stall scores at 30 min associated with each structural class and in function of STR length. The reported structures are those predicted when the STRs are 72 nt long. Centre lines denote medians, boxes span the interquartile range, and whiskers extend beyond the box limits by 1.5 times the interquartile range. *P* values for the comparison of the distributions were calculated using Kolmogorov–Smirnov tests, n.s *P* > 0.05, \**P* ≤ 0.05, \*\**P* < 0.01, \*\*\**P* < 0.001. **(f)** Hierarchical clustering of STR single-stranded motifs by sequences using cosine distances as measure of sequence similarity together with heatmaps reporting GC content and sequence entropy. The leaves of the dendrogram are coloured according to the predicted structures of the STRs. Such representation highlights two distinct families related to G4 (red) and i-motifs (green) forming STRs. Hairpin-like STRs are dispersed over the dendrogram showing a higher degree of sequence diversity.

We used ^1^H NMR spectroscopy to validate some of the predicted structures (Additional file 1: Figure S5). Of specific interest, we confirmed that both the G and C mononucleotide repeats were folded under the condition of our assay into structures consistent with a G4 and i-motif respectively. We also validated the structure of STRs predicted to fold into the more elusive i-motifs showing that the hexanucleotides repeats ATCCTC and ACCCGG both fold into structures consistent with i-motifs at physiological pH (pH 7.5). Finally, we found that, unexpectedly, the GA dinucleotide repeat can fold into a G4 structure under the condition of our assay. Taken together these results demonstrate that our assay can infer STR structure from polymerase stalling events.

### Hairpin-like and tetrahelical structures arise from distinct STR families with distinct properties

We next investigated the different STR structural classes further. Interestingly, while the three structural classes displayed similar stall scores at 0.5 min (Fig. 3b) their stall scores differ significantly at 30 min (Fig. 3c) with the QUAD displaying higher scores than the IMOT and HAIRP sequences, confirming that while STRs predicted to fold into G4s induce persistent DNA polymerase stalling, stalling events at hairpins and i-motifs are resolved over time by the polymerase. As expected, the longer an STR, the more potently it was able to induce stalling. Indeed, while 17.4% of the STR are predicted to be folded when 24 nt long, this fraction increases to 30.7 and 37.5 % when they are 48 and 72 nt long respectively (Fig. 3d). We then interrogated the association between the periodicity of STRs and their ability to stall polymerase progression. We found that longer structured STR motifs, *i*.*e*. those with longer periodicity, are associated with lower stall scores (Additional file 1: Figure S4c) and that the number of STR motifs correlated with the stall scores (Additional file 1: Figure S4d). These observations suggest that shorter STR motifs, *i*.*e*. shorter periodicity, may have more base pairing opportunities leading to more stable DNA structures. Interestingly, the stall scores of short 24 nt HAIRP repeats at 30 min were not statistically different from the control sequences (Fig. 3e) indicating that STRs folding into hairpins need to be relatively long to induce persistent polymerase stalling. On the other hand, stalling at STRs folding into tetrahelical structures is persistent even for short 24 nt repeats (Fig. 3e). This observation suggests that STRs of different structural classes might be expected to evolve differently within genomes, a point to which we return below.

To assess the extent to which structure-forming STRs share common sequence features, we created a dendrogram reporting the hierarchical sequence relationship between STR sequences (see Methods details) and highlighted their structural class (Fig. 3f). This clustering representation distinguishes the two main families of tetrahelical STRs, while hairpin-like STRs are scattered throughout the dendrogram showing that they exhibit a greater degree of sequence diversity than tetrahelical STRs. This conclusion is further supported by the observation that while the sequences of HAIRP STRs are characterised by a higher entropy but a similar GC content to UNF STRs, the sequences of QUAD and IMOT STRs are characterised by a lower entropy and a higher GC content than the UNF STRs (Additional file 1: Figure S4e and S4f).

We then interrogated the nature of hairpin-like STRs to assess the sequence requirement for hairpin formation. We found that only 37 out of the 188 (∼ 20%) unique single-stranded STR motifs predicted to fold into hairpin-like structures are palindromic sequences suggesting that the majority of motifs lead to imperfect hairpin, *i*. *e*. stem loop structures with mismatches. We used the Mfold DNA folding algorithm [30] to predict the most stable hairpins arising from these motifs, identify potential mismatches and characterise the nature of the bases involved in base pairing. We found that the longer an STR, the more mismatches are tolerated (Additional file 1: Figure S4g) and that increased number of mismatches are associated with lower stall scores (Additional file 1: Figure S4h). In this context, hairpin-like structures are enriched in G•C base pairings (odds ratio = 1.82; Fisher’s two-sided P = 1.21 x 10^−4^) and mismatches involving As (odds ratio = 1.60; Fisher’s two-sided P = 1.42 x 10-3). The latter point suggests that A may stabilise mismatches and/or are involved in non Watson-Crick base interactions. For example, we found that hairpin-like structures arising from the (CAG•CTG) repeat display different stall scores (Additional file 1: Figure S4i) with higher scores for the CAG strand displaying A-A mismatches that for the CTG strand displaying T-T mismatch.

### The DNA polymerase remodels STRs during DNA synthesis

Our approach for recovering stalled DNA synthesis products allows us to map, at single nucleotide resolution, the last nucleotide incorporated during the primer extension reaction and therefore the site of polymerase stalling (Fig. 4a and Additional file 1: Figure S6a). By computing the mean distance of the stalling sites from the start of the repeat for each sequence, we found that the mapped position of the stalled DNA polymerase is time dependent with increasing values over time (Additional file 1: Figure S6b) suggesting that, even in the event of polymerase stalling, DNA synthesis can occur within the repeats. We found that the distance travelled by the DNA polymerase within a repeat directly anticorrelates with its stall score (Fig. 4b). For example, while the DNA polymerase is able to progress on average by 17 nt, and up to 65 nt, within repeats with low σ, DNA synthesis within repeats with a high σ does not progress, on average, further than 2 nt. The distance travelled within a repeat also depends on the structural class of the STRs, with the DNA polymerase able to travel further through hairpin-like STRs than tetrahelical STRs (Fig. 4c). This observation suggests that the DNA polymerase can, in some circumstances, replicate through STRs by remodelling the structures formed on the template even without any accessory factors such as helicases or single strand binding proteins. Hence the polymerase can, possibly through successive rounds of dissociation/re-association [31], overcome DNA structure and resume replication in some circumstances. Taken together these results suggest that DNA synthesis at structured STR is a highly dynamic process in which the elongation of the DNA template can remodel STR secondary structure.

**Fig. 4.**
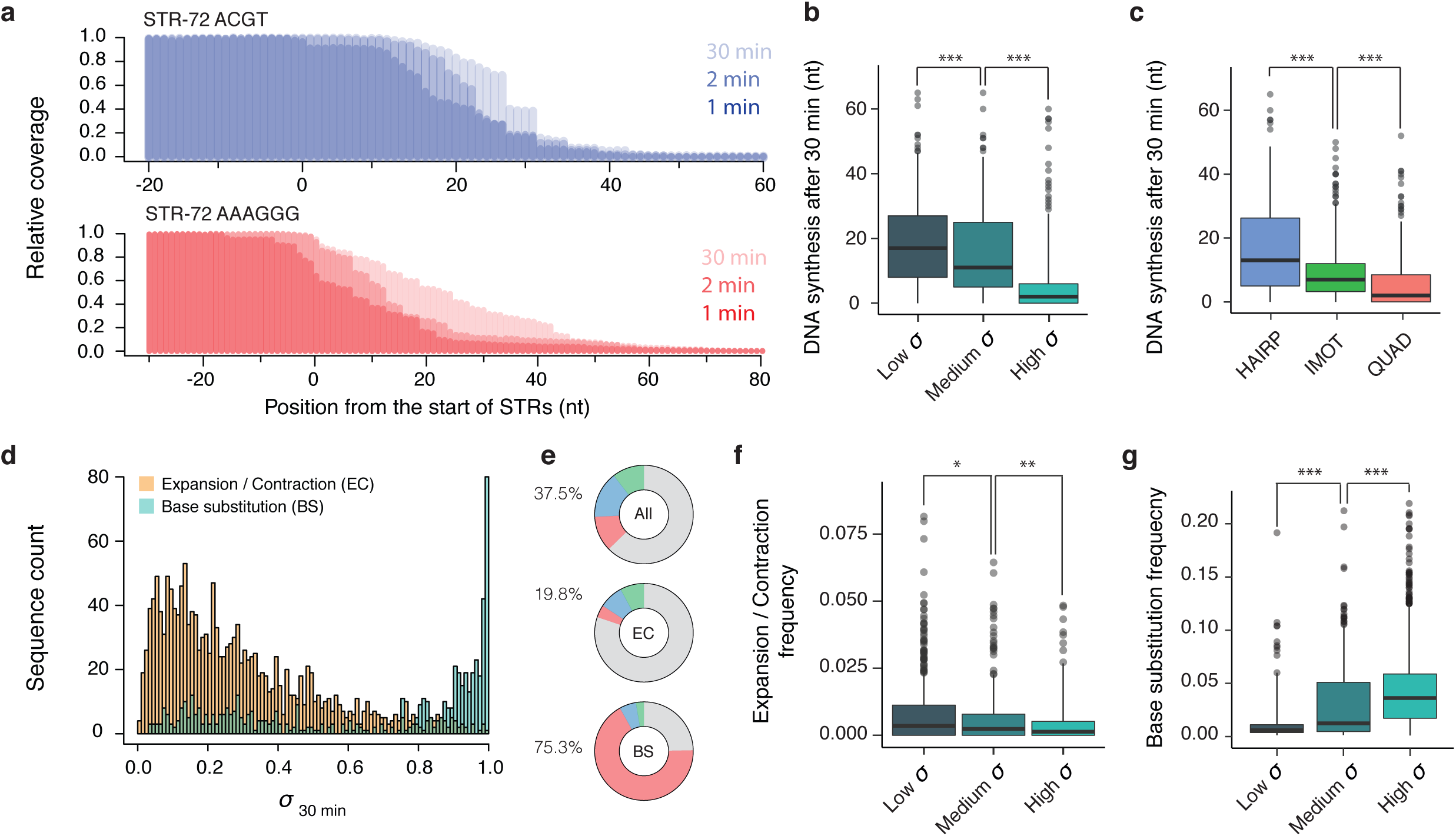
DNA synthesis at structured STRs is a highly dynamic and error-prone process. Deep sequencing of the stalled products of replication allows DNA synthesis to be followed over time and the mapping of stalled DNA polymerase at single nucleotide resolution. **(a)** Plotting the relative reads coverage at STRs, *e*.*g*. the 72 nt long ACGT and AAGGG repeats folding into hairpin-like and G4 structures respectively, reveals time-dependent sharp transitions corresponding to the main positions where the DNA polymerase stalls and dissociates during DNA synthesis. Computing the distance of the stalling sites, *i*.*e*. the mid-transition value, from the start of the repeat for each STR at different time points shows that the distance travelled by the polymerase after 30 min depends on **(b)** the stall scores of the STR and **(c)** their structures. 72 nt long STRs were binned either according to low (σ < 0.33), medium (0.33 ≤ σ < 0.66) and high (σ ≥ 0.66) stall scores at 30 min in **(b)** or to their predicted structures in **(c)**. Deep sequencing of the extended products of replication allows the identification of new sequence variants within the pool of newly synthesised DNA molecules. Mutations were classified into either base substitutions (BS) or expansion/contraction events (EC). **(d)** Distribution of the stall scores at 30 min associated to STRs presenting either expansion/contraction (orange) or base substitutions (light blue) events. **(e)** Pie charts representing the proportion of STR folding into hairpin-like (blue), i-motifs (green) or G4s structures (red) or unfolded (grey) when considering all STRs (All), STRs marked by expansion/contraction (EC) or base substitutions (BS) events. These charts show a depletion and an enrichment of structured STRs within the pool of STRs characterised by presenting either length or sequence variation respectively. Frequencies of **(f)** expansion/contraction and **(g)** base substitution events observed at STRs when binned according to their stall scores at 30 min (low: σ < 0.33, medium: 0.33 ≤ σ < 0.66, high: σ ≥ 0.66). Frequencies are defined as the ratios of the number of reads supporting a mutation by the total number of reads covering this mutation. Centre lines denote medians, boxes span the interquartile range, and whiskers extend beyond the box limits by 1.5 times the interquartile range. *P* values for the comparison of the distributions were calculated using Kolmogorov–Smirnov tests, \**P* ≤ 0.05, \*\**P* < 0.01, \*\*\**P* < 0.001.

### DNA polymerase stalling triggers sequence instability

We then assessed the extent to which STRs influence the fidelity of DNA synthesis by identifying new sequence variants generated during the primer extension reaction (see Methods). We classified the sequence variants into either point mutations or polymerase slippage events (expansions or contractions). We identified 644 unique point mutations and 1740 slippage events distributed among 250 and 618 unique single-stranded STR motifs respectively. Examples visualising STR alignment and mutation called are shown in Additional file 1: Figure S7. We combined all sequence variants identified at any time point of our extension assay and determined the correlation between the stall scores of the reference STR motifs and their stability (Fig. 4d). Surprisingly, we found that while STRs with low stall scores are more prone to expansion/contraction, STRs of high stall scores are more prone to point mutation, suggesting STR structural class influences the pattern of mutagenesis in our primer-extension assay. Supporting this observation, we found that structured STRs are enriched within the pool of STRs exhibiting point mutation (odds ratio = 5.10; Fisher’s two-sided *P* < 2.2 × 10^−16^) and depleted (odds ratio = 0.41; Fisher’s two-sided *P* < 2.2 × 10^−16^) in those exhibiting length variation (Fig. 4e).

These observations suggest that the absence of structure within STRs globally favours their ability to expand and contract during DNA synthesis. Supporting this observation, we found that the frequency of slippage events anticorrelates with the stall scores of the STRs (Fig. 4f). Among the structured STRs we found that those folding into hairpin-like and i-motifs were the most frequently expanded or contracted (*P* ≤ 6.5 x 10^−14^, Additional file 1: Figure S8a). We then analysed the slippage events to assess whether structures affect the step-size distribution of the slip. While the majority of observed events were gain of single repeat unit, we found that HAIRP STRs are more likely to mutate by multiple units at once and are more prone to contraction (odds ratio = 3.42; Fisher’s two-sided *P* = 1.39 × 10^−5^, Additional file 1: Figure S8b). Taken together these observations show that while structured STRs are more stable than their unfolded relatives in this experiment, the nature of their structure affects their slippage mutation pattern and suggest that STR length within the genome may evolve differently according to their structure.

We next investigated the relationship between point mutation frequency and STR structure. We found that STRs with high stall scores are more frequently mutated (Fig. 4g), interrupting their repetitive sequence, and that the number of base substitutions per repeat increases with the ability of the STR to stall polymerase (Additional file 1: Figure S9a). Interestingly, point mutations increase in frequency between the start and end of the repeats (Additional file 1: Figure S9b) indicating that the chances of misincorporation by the polymerase increase as a function of the distance synthesised through the repeat. Among the structured STRs, the QUAD class displays higher mutation rates with point mutations up to 5.4-fold more frequent than in HAIRP and IMOT STRs (Additional file 1: Figure S9c). This observation shows that polymerase fidelity is affected in distinct ways by different classes of DNA structure. We then computed the frequency of each base substitution event (Additional file 1: Figure S9d) and assessed their representation among each STR structural class (Additional file 1: Figure S9e and S9f). Surprisingly, while the overall pattern of substitutions is similar in the UNF STRs and control sequences, each structural class displays a unique mutational signature (Additional file 1: Figure S10a). These structure-specific mutational signatures could not be explained by biases in base composition (Additional file 1: Figure S10b) and were found to be due to the T7 DNA polymerase (Additional file 1: Figure S10c). Interestingly, QUAD STRs were mainly mutated at non-guanine nucleotides, *i*.*e*. bases within spacer sequences that do not contribute to the stabilisation of the G4 motif, which is consistent with a recent analysis of mutation of G4 motifs in cancer genomes [32]. Taken together these observations suggest that STR length and sequence instability is intimately linked to polymerase stalling due to STR structure formation.

### DNA polymerase stalling at DNA structures predicts STR abundance and length in eukaryotic genomes

We next aimed to harness the information from our high-throughput assay to examine the impact of DNA polymerase stalling at STRs on their genomic representation. Using data from the MicroSatellite DataBase [3] reporting ∼ 4,500,000 STR loci within the human genome, we found that the relative abundance of each of the 501 unique double-stranded motifs directly anticorrelates with the ability of the motif to stall DNA polymerase (Fig. 5a) suggesting that STRs capable of significant secondary structure are deleterious. Indeed, structured STRs are less abundant that their unstructured counterparts (*P* ≤ 0.009615, Additional file 1: Figure S11a). Interestingly, among the different structural classes, the HAIRP STRs were least abundant (Additional file 1: Figure S11a). These observations suggest that even transient DNA polymerase stalling in STR loci poses a challenge to replication and can trigger genomic instability. We postulated that deleterious STRs may be maintained at minimal length in order to minimise their impact. We found that STRs with high stall scores are statistically shorter than STRs with low stall scores (*P* < 2.2 x 10^−16^, Additional file 1: Figure S11b and S11c) and that the genomic length range of STR motifs decreases as their stall score increases (Fig. 5b). For example, while the human genome can accommodate long low stall score STRs (*e*.*g*. the AACCCT motif can be as long as 1103nt), high stall score STRs are maintained at short length (*e*.*g*. the ATCCGG motif does not exceed 16 nt). We found that these global trends are largely conserved within the genome of five other eukaryotic species such as *Mus musculus, Gallus gallus, Danio rerio, Drosophila melanogaster* and *Saccharomyces cerevisiae* (Additional file 1: Figure S12). The most abundant STR within these species is the AAAT motif, which was found to be unfolded under the conditions of our assay, while the least abundant are the ACGCGT, ATCGCG and ACGTCG motifs predicted to fold into hairpin-like structures displaying stall scores above 0.8 at 0.5 min when 48 nt long. These observations suggest that structured STRs impose a replicative burden, which hampers their maintenance in eukaryotic genomes.

**Fig. 5.**
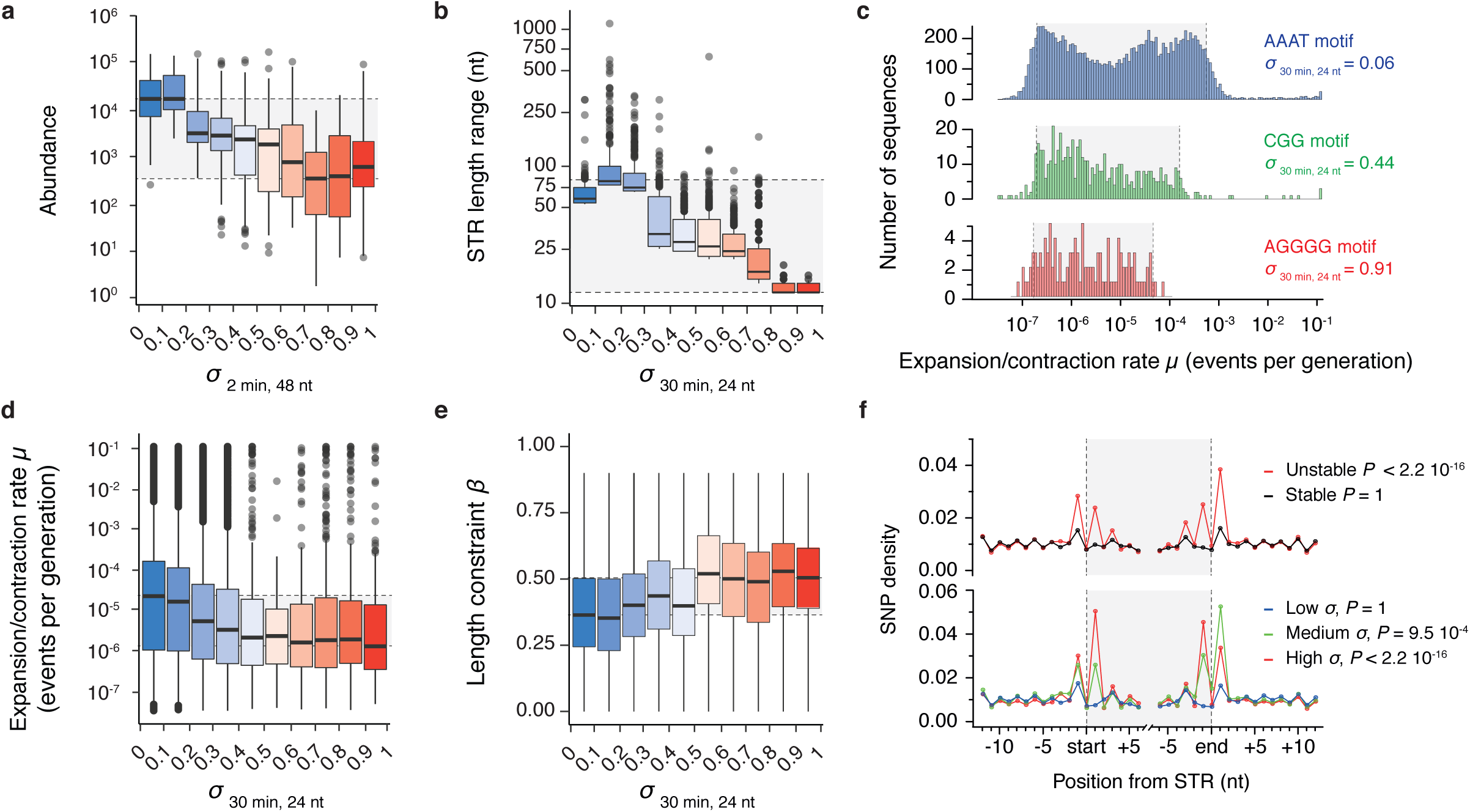
DNA polymerase stalling at DNA structures predicts abundance, length and stability of STRs in the human genome. **(a)** Abundance, *i*.*e*. number of occurrences, of the 501 unique double-stranded STR motifs in the human genome when binned by their average stall scores at 2 min when 48 nt long. **(b)** Range of double-stranded STR motifs length when binned by their average stall scores at 30 min when 24 nt long. The plot reports the top 1,000 longest repeat instances from each bin. Similar analyses related to other eukaryotic genomes are reported in Additional file 1: Figure S12. **(c)** Distribution of expansion/contraction rates of the AAAT, CGG and AGGGG motifs which are characterised by low, medium and high stall scores at 30 min when 24 nt long respectively. Shaded boxes span from the 5th to the 95th percentiles of the distributions. **(d)** Expansion/contraction rates (*μ*) and **(e)** length constraints (*β*) associated with the double-stranded STR motifs when binned by their average stall scores at 30 min when 24 nt long. The plots show a correlation between the length stability and the ability of a STR to stall polymerase, which is due to an increased length constraint reflected by higher β values. **(f)** SNP density at positions surrounding STRs when binned according to their stability (top) or stall scores at 30 min when 24 nt long (bottom, low: σ < 0.33, medium: 0.33 ≤ σ < 0.66, high: σ ≥ 0.66). SNP density at individual positions was computed by dividing the number of STR marked by a SNP at a given position by the total number of considered STRs. Sawtooth profiles in SNP density is due to the underlying periodicity of the repeats (see Additional file 1: Figure S14a). Centre lines denote medians, boxes span the interquartile range, and whiskers extend beyond the box limits by 1.5 times the interquartile range. Shaded boxes highlight the range of the medians. Reported *P* values assess SNP enrichment in the vicinity of STR and was computed using two-sided Fisher tests considering the average frequency of SNPs at position −1 and +1 from the start and the end of the repeats.

### The lengths of structured STRs are constrained and stable within the human genome

In order to assess how structured STRs and their unfolded counterparts evolve within the human genome, we took advantage of a recent dataset that provides an estimate of mutation parameters for ∼ 1,250,000 STR loci in the human genome [24]. In this work the authors develop a model that quantifies three parameters for each STR locus: a per-generation length mutation rate (μ), a length constraint (*β*) and a mutation step size distribution (*p*). *μ* represents the rate of expansion and contraction observed at a given STR locus, *β* quantifies the evolutionary force that constrains a given STR to its reference length and *p* denotes the probability that a mutation occurs at a single STR unit at a time. The combination of these three parameters allows modelling of the evolutionary pathway of each human STR and describes their stability over time.

We assessed the correlation between each parameter and the stall score of each STR locus (see Methods). Because polymerase stalling is associated with genetic instability, we were expecting a direct correlation between the stall score of a STR motif and its rate of length variation, but found the opposite. For example, the STR motif showing the highest average rate of expansion/contraction is the low stall score AAAT motif with a median mutation rate of 1.3 x 10^−5^ events per generation while the high stall score AGGGG motif exhibited a median mutation rate of 1.1 x 10^−6^ events per generation (Fig. 5c). Analysis of the 501 unique double-stranded motifs showed that higher stall scores were associated with less length variation (Fig. 5d) suggesting that the length of structured STRs is stable over time. Among the structured STRs we found that the tetrahelical STRs show the highest length stability and estimated that the unfolded STRs are ∼12 times more likely to expand or contract (Additional file 1: Figure S13a). We postulated that the lower length variation observed for structured STRs may be due to greater length constraints. Indeed, length constraints (*β*) associated with high stall score STRs are ∼1.5 time higher than those associated with low stall scores STRs (Fig. 5e). Among the structured STRs, the hairpin-like STRs are the most constrained sequences (Additional file 1: Figure S13b). Taken together these results show that the length of structured STRs is more constrained than their unstructured counterparts.

We then investigated the impact of structures within STRs on the distribution of the expansion / contraction step size, *p*. We found that the higher the stall score of an STR, the higher the probability that it expands or contracts by more than one unit at a time (Additional file 1: Figure S13c). Among the structured STRs, the HAIRP class displays the greatest propensity to mutate by more than one unit at a time (Additional file 1: Figure S13d), consistent with the results from our primer extension assay. Together, these results suggest that DNA polymerase stalling at structured STRs is a determinant of length stability.

### Structured STRs are prone to point mutation in the human genome

The greater sequence instability of structured STRs in our primer extension assay (Fig. 4d and 4g) suggested that increased length constraint of structured STRs within the human population may be driven by point mutagenesis disrupting the repeat pattern. To test this hypothesis, we leveraged population scale genomic data from the NCBI database [33] reporting ∼34 million germline mutations and asked whether SNPs are enriched in the vicinity of STR loci. We initially assessed SNP density at positions surrounding STRs of stable or unstable length, *i*.*e*. STRs with length mutation rates below and above 10^−7.5^ events per generation respectively. We found that while positions surrounding STRs of stable length are characterised by a basal level of SNPs, the positions directly upstream and downstream from the start and end of STRs of unstable length are enriched in SNPs (odds ratio = 2.53; Fisher’s two-sided *P* < 2.2 × 10^−16^, Fig. 5f). We then binned the STRs of unstable length according to their stall scores and found that the higher the stall score of a STR, the higher the density of SNPs at the boundary of the locus (Fig. 5f). We found that the enrichment at high stall score STRs (odds ratio = 2.18; Fisher’s two-sided *P* < 2.2 × 10^−16^ for σ _30 min, 24 nt_ ≥ 0.66) is more pronounced than at medium stall score STRs (odds ratio = 1.08; Fisher’s two-sided *P* = 0.00095 for 0.33 ≤ σ _30 min, 24 nt_ < 0.66). In fact, SNP densities are higher at position surrounding high stall score than medium stall score STRs (Fisher’s two-sided *P* < 2.2 × 10^−16^). This observation suggests that DNA polymerase stalling at structured STR induces error-prone DNA synthesis, which is consistent with the results from our primer extension assay. Indeed, we found that hairpin-like and tetrahelical STRs are marked by SNPs at their boundaries (Additional file 1: Figure S14b). Interestingly, we found that each structural class is characterised by a specific and unique substitution matrix (Additional file 1: Figure S14c) that could not be explained by biases in base composition (Additional file 1: Figure S14d). Finally, we found that while the boundaries of long STRs are depleted in SNPs (odds ratio = 0.84; Fisher’s two-sided *P* = 0.0173 for *Length* > 64 nt), the boundaries of short STRs are enriched in SNPs (odds ratio = 2.98; Fisher’s two-sided *P* < 2.2 × 10^−16^ for *Length* ≤ 18 nt, Additional file 1: Figure S14e). Taken together with the increased rates of base substitution at structured STRs observed in our primer extension assay, these data suggest that error-prone DNA synthesis at DNA structures constrains their expansion.

## Discussion

We present in this study a high-throughput primer extension assay for measuring the kinetics of DNA synthesis and DNA polymerase stalling at all STR permutations of different lengths. Using a model replicative polymerase in a deliberately minimalist system representing the most fundamental aspect of DNA replication, *i*.*e*. templated DNA synthesis in the absence of accessory factors, we demonstrate that DNA polymerase stalling at structured DNA is sufficient to describe complex features of eukaryotic genomes such as STR abundance, length and stability. Our results do not exclude other mechanisms that contribute to STR length variation, *e*.*g*. various DNA repair pathways, but highlight the central role of replicative DNA synthesis in sculpting STRs within the genome. Our assay takes advantage of a polymerase that has been engineered to lack proofreading activity, allowing normally rare mutational events to be detected within the course of a single round of DNA synthesis. It is noteworthy that even if the T7 DNA polymerase, used in our assay, differs from the B-family of eukaryotic replicative polymerases, its behaviour at structured STRs allows understanding their evolution in eukaryotic genomes. This observation support that DNA structures are the main determinants of STR stability.

Our approach has uncovered the nature and extent of STRs that impede DNA synthesis. We identified unanticipated DNA structures within simple repeats, for instance the formation of i-motifs within the C mononucleotide repeat and quadruplexes by the GA dinucleotide. Indeed, structures were formed that were thermodynamically stable enough to stall polymerase under the condition of our assay by 37.5 % of the STRs. However, we anticipate that other repeats could fold into metastable structures *in vivo* that may interfere with other biological processes such as nucleosome positioning or transcription factor binding. Our data hence represent a useful resource for discovering structures arising from low complexity sequences.

By leveraging population-scale genomic data, we found that structure formation by STRs is associated with greater length stability but increased single-nucleotide mutation rates in the human genome. Our observations support DNA polymerase stalling at DNA structures at STR loci being a key determinant of both length and sequence mutation rates and explains the selective length constraint of structured STRs. Overall, our data support a model in which the length of structure-prone STRs at evolutionary equilibrium results from a balance between expansion and point mutations regulated by polymerase stalling at DNA structures (Fig. 6). After initiation of DNA synthesis, if the DNA polymerase dissociates within an STR, its repetitive nature may induce an out-of-register realignment of the newly synthesised DNA on the template strand, leading to the expansion of the repeat according to a strand slippage mechanism [34]. This process may occur several times until the STR reaches an equilibrium length. While the equilibrium length of an unstructured STR will be defined by the difference between the length-dependent expansion and contraction rates [21], the maximum length of a structured STR is defined by its ability to stall polymerase progression. Indeed, structured STRs may expand until reaching a threshold length at which the repeat will accommodate thermodynamically stable structures that may stall DNA polymerase and trigger error-prone DNA synthesis. Error-prone DNA synthesis at structured STRs is supported by both our data from our primer-extension assay and the observation of increased densities of SNPs at the boundaries of STRs of high stall scores within the human genome. This observation is also in line with a recent work reporting a concurrent change in repeat length with mutagenesis trigger by replication fork stalling at the Fragile X CGG repeats in mammalian cells [35] and the observation of higher rates of nucleotide mutation at non-B DNA regions of the human genome [36]. Sequence variation at the boundaries of the structured STRs will then dilute the repetitive sequences within the flanks of the STR loci. The length of the structured repeat at equilibrium is then a result of a balance of the two processes of strand slippage and error-prone DNA synthesis. Because the stability of DNA structures and the ability of structured STRs to stall DNA polymerase increase with the length of the repeats, this model explains why structured STRs are kept at minimal lengths. DNA polymerase stalling will also influence which STRs will be maintained or spread which is reflected by a decreased abundance of structured STRs. Hence error-prone DNA synthesis at structured DNA can be seen as a driver of purifying selection for deleterious STR.

**Fig. 6.**
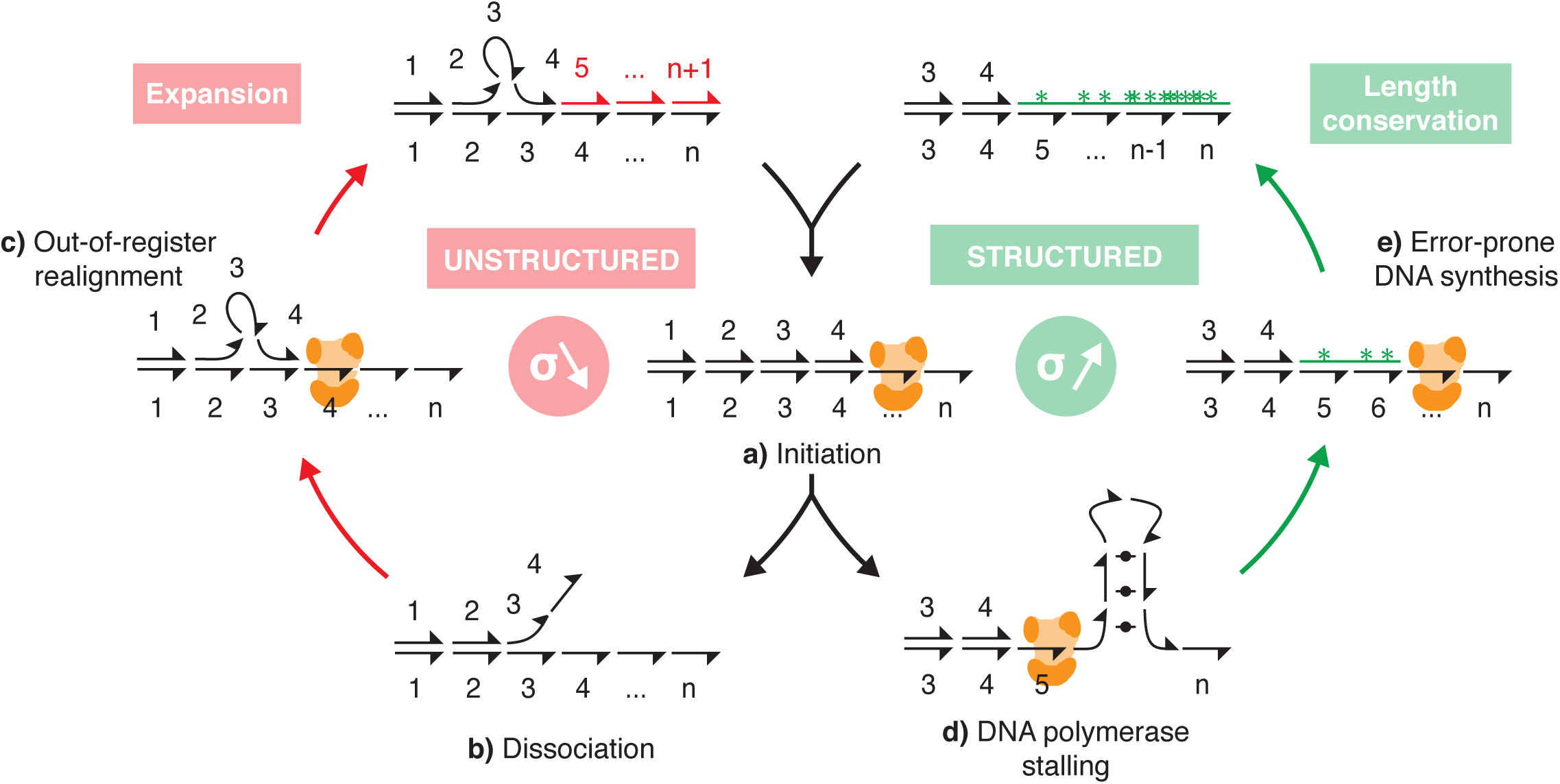
A model for the regulation of the length of structured STRs. After initiation of DNA synthesis **(a)**, if the DNA polymerase dissociates within an STR **(b)**, its repetitive nature may induce an out-of-register realignment **(c)** of the newly synthesised DNA on the template strand, leading to the expansion of the repeat according to a strand slippage mechanism. After several rounds of expansion, the STR may reach a threshold length at which the repeat will accommodate thermodynamically stable structures that would trigger polymerase stalling **(d)** and induce error-prone DNA synthesis **(e)**. Base substitution at the boundary of the structured STRs then dilute the repetitive sequences within the flanks of the STR locus. The length of the structured repeats at equilibrium therefore results from the balance of two processes: strand slippage and error-prone DNA synthesis regulated by polymerase stalling at DNA structures.

## Conclusions

Our work not only provides a comprehensive characterisation of DNA synthesis at all STR permutations but also unravels the interplay between STR structures and genomic STR stability. We have analysed general trends associated with structured STRs but we note that some repeats, particularly those associated with neurological disorders, do not seem to be similarly constrained and can expand to lengths that are detrimental to the physiology of the cell. This suggests that additional forces operate to drive pathological expansion beyond the length constraints suggested by this study. Overall, our work provides a valuable dataset for interpreting the evolution of structured STRs in eukaryotic genomes and reveals a previous unappreciated role for polymerase behaviour at DNA structures in genome stability and evolution.

## Methods

### Library design

We designed a total of 20,000 sequences, divided into several structural classes, each aimed at examining the impact of DNA structure on polymerase stalling (Additional file 2). As positive controls, *i*.*e*. sequences designed to fold into known single-stranded DNA structures, we designed 960, 1500 and 472 sequences folding into hairpins, G-quadruplexes and i-motifs respectively. Hairpins are sequences (7 to 22 nt) of varying GC content (from 20 to 80%) with their reverse complement downstream a T4 linker. G-quadruplexes are sequences composed of four G-tracts (2 to 5 nt) separated by random sampling of loops (1 to 9 nt). i-motifs are sequences composed of four C-tracts (2 to 5 nt) separated by random sampling of loops (1 to 9 nt). This randomised approach allows building a library of sequences folding into structures of different stabilities. To control for DNA stalling due to extreme GC values, we incorporated to the library 1,000 72 nt long random sequences of varying GC content (from 20 to 80%) as negative controls. The STR sequences cover all 5,356 possible permutations of 1 to 6 nt motifs and are present in the library in three different length (24, 48 and 72 nt). The 3’ edge of each sequence contains a unique 12 nt sequence, which was used as a barcode that allowed to uniquely identify the associated sequence within the sequencing reads. Every barcode differs from any other barcode in at least 3 nt excluding low complexity sequences (homopolymers longer or equal at 3 nt), allowing the correct identification of any sequences even if it contains a single-base mutation. Common priming sites and restriction sites at both ends flank each sequence. The exact sequences composing the library are reported in Additional file 2. The library was synthesised on a programmable microarray by Twist Bioscience and received as a pool of 20,000 different single-stranded 150 nt long oligonucleotides.

### Preparation of the designed sequence library

The pool of oligonucelotides was dissolved in 10 mM Tris.EDTA pH 7.4 at a final concentration of 10 ng/μL. 20 ng of the single-stranded library DNA was split into 20 PCR amplification reactions in a final volume of 25 μL. Each reaction contained 0.75 μL of 10 μM forward and reverse primer mix, 5 % DMSO and 12.5 μL of KAPA HiFi HotStart ReadyMix (Roche). The primers used to amplify the library were P1 (5’-GGGGAAGCTTGCCGTAAG-3’, forward primer) and P2 (5’-TGATCGCGGATCCATCGC-3’, reverse primer). The parameters for PCR were 95 °C for 5 min, 10 cycles of 98 °C for 20 s, 60 °C for 15 s and 72 °C for 30 s and then one cycle of 72 °C for 30 min. The PCR products from all reactions were pooled and purified with the Monarch PCR & DNA Cleanup Kit (NEB) according to the manufacturer’s instructions.

### Ligation and transformation

Purified DNA library (250 ng) was cut with the Hind III and Bam HI restriction enzymes (NEB) for 1h at 37 °C in a reaction mixture containing 1X of the NEB cut smart buffer. Digested DNA was purified with the Monarch PCR & DNA Cleanup Kit (NEB) according to the manufacturer’s instructions. To prepare the vector for cloning and amplification, 1.5 μg of the pBluescript II SK (–) phagemid (Agilent) was cut with Hind III and Bam HI (NEB) for 1h at 37 °C, treated with alkaline phosphatase (Roche) and purified with the Monarch PCR & DNA Cleanup Kit (NEB) according to the manufacturer’s instructions. The DNA library was then ligated to the linearised pBluescript II SK (–) plasmid using T4 DNA ligase (NEB) at a 3:1 insert to vector ratio using 200 ng of the linearised vector. The ligation product was purified using the Monarch PCR & DNA Cleanup Kit (NEB) eluting the product in 10 μL. Ligated DNA was transformed into four tubes, each containing 25 μL of *E*. *coli* XL1-Blue electroporation-competent cells (Agilent), which were then plated on 6 25-cm agar plates containing 2xTY medium broth and ampicillin. Sixteen hours after transformation at 37 °C, the plates contained ∼ 8,000 colonies per cm^2^ which represent a ∼ 1500 X coverage of the DNA library. Each plate was scraped into 2xTY medium containing ampicillin and the cells were incubated for 30 min at 37 °C in a total volume of 400 mL. This cell suspension was used to prepare 1 mL glycerol stocks (85% cell suspension, 15 % glycerol) that were freeze on dry ice and store at −80 °C for later use.

### Preparation of the single-stranded DNA template

Glycerol stocks from the previous step were thawed on ice and expanded in 50 mL of 2xTY medium at 37 °C for 1h. Cell cultures were then infected with 50 μL of M13KO7 helper phage (final concentration at 1.10^8^ pfu/mL) and incubated at 37 °C for 1 h. Kanamycin was added at a final concentration of 70 μg/mL and the cultures were grown overnight at 37 °C shaking at 250 rpm. *E*. *coli* cells were removed from the solutions by two centrifugations at 4,000 g for 10 minutes. 90% of the supernatant was transferred into a new tube and 0.2 volume of a 2.5 M NaCl, 20% PEG-8000 solution was added. The resulting solutions were gently mixed and incubated at 4 °C for 60 minutes. Phage particles were recovered by centrifugation at 12,000 g for 10 minutes. Phage pellets were resuspended in 1.6 ml TBS and spun in a microfuge for 1 min (2500 rpm). 800 μL of the supernatants were transferred to new tubes and 160 μL of the 2.5 M NaCl, 20% PEG-8000 solution was added. The solutions were incubated at room temperature for 5 min, spun in a microfuge for 10 minutes at high speed. Each phage pellet was resuspended in 300 μl 10 mM Tris.EDTA pH 7.4 and extracted once with phenol, then twice with phenol/chloroform (50/50: v/v), and finally chloroform. The single-stranded DNA was recovered by ethanol precipitation and resuspended in 50 μL of 10 mM Tris.EDTA pH 7.4. Each preparation allows the recovery of ∼ 50 μg of the single-stranded DNA library. A similar protocol was used to prepare single-stranded template containing individual sequences such as a G4 (TGGGAGGGTGGGAGGG) or a mutated G4 (GGGA<u>CCC</u>TGGGAGGG).

### High-throughput primer extension assay

Primer extension reactions were performed using a modified T7 DNA polymerase (Sequenase™ Version 2.0, Thermofisher) and a Cy5-labeled primer (P3: 5’-Cy5-TAATGTGAGTTAGCT-3’) annealing to the 3098 nt ssDNA templates and 230 nt from the start of the STRs or designed structures. Each reaction contained 0.5 μL of 10 μM fluorescently labelled P3, 2.5 μL of 100 ng/μL of the ssDNA templates in 7 μL of the reaction buffer. The final composition of the reaction buffer was 40 mM Tris.HCl pH 7.5, 20 mM MgCl_2_, 50 mM NaCl and 50 mM KCl. The mix was incubated for 2 min at 80 °C and slowly cooled down to 20 °C over 1 h. 1 μL of 0.1 M DTT, 1.5 μL of 10 mM dNTPs and T7 DNA polymerase (1.625 units) were then added for a total volume of 15 μL. The reactions were carried out at room temperature. At the indicated time points, the primer extension reactions were stopped by adding a formamide loading dye (95 % formamide, 20 mM EDTA, 0.05 % bromophenol blue, 0.05 % xylene cyanol) and the products were separated on a 6 % urea-PAGE gels (Invitrogen) run at 180 V for 50 min. To assess the position of the stalled products, the assay was performed using two templates containing either a G4 motif (GGGAGGGTGGGAGGG) or a mutated form of the same sequence (GGGA<u>CCC</u>TGGGAGGG) not expected to support G4 formation.

### Sample preparation for sequencing

The bands corresponding to the fully extended and stalled products were excised using the products of replication from the G4 or mutated G4 containing templates as guide. Because the stalled products associated with STR sequences are expected to run as a smear, the band corresponding to the stalled products was excised from the bottom of the corresponding band to the first noticeable transient pause sites. The gel matrix was then crushed, soaked in 400 μL of 500 mM sodium acetate supplemented with 1 μL of 20 % SDS. The samples were rocked at 37 °C for 2 h, filtered in Costar Spin-X centrifuge tube filters (Sigma) and DNA was recovered by ethanol precipitation. Pellets were resuspended in 30 μL of 10 mM Tris.EDTA pH 7.4. Stalled products were then prepared for PCR amplification by adding poly-dC homopolymer tails to their 3’ ends using a terminal transferase (TdT). TdT tailing was performed using 10 μL of the recovered DNA in a final volume of 50 μL containing 5 μL of 2.5 mM CoCl_2_, 2.5 μL of 2mM dCTP, 20 units of TdT (NEB) and the provided TdT buffer. The reactions were incubated at 37 °C for 30 min and heat inactivated at 70 °C for 10 min. Each sample was split into 5 PCR amplification reactions in a final volume of 25 μL. Each reaction contained 0.75 μL of 10 μM P2 primer, 0.75 μL of 10 μM anchor primer (5’-GGCCACGCGTCGACTAGTACGGGIIGGGIIGGGIIG-3’ where I is inosine), 5 % DMSO and 12.5 μL of KAPA HiFi HotStart ReadyMix (Roche). The parameters for PCR were 95 °C for 5 min, 35 cycles of 95 °C for 15 s, 53 °C for 30 s and 72 °C for 30 s and then one cycle of 72 °C for 5 min. The PCR products from all reactions were pooled for purification. PCR amplifications of the extended products and the parental library, *i*.*e*. the library that hasn’t been extended by the T7 DNA polymerase, were performed using the exact same conditions in order not to introduce PCR biases due to the presence of structures within the templates. 5 μL of the recovered DNA and 1 ng of the parental library were amplified in reactions of 25 μL. Each reaction contained 0.75 μL of 10 μM P1 and P2 primer mix, 5 % DMSO and 12.5 μL of KAPA HiFi HotStart ReadyMix (Roche). The parameters for PCR were 95 °C for 5 min, 10 cycles of 98 °C for 20 s, 60 °C for 15 s and 72 °C for 30 s and then one cycle of 72 °C for 30 min. The PCR products from all reactions were purified with the Monarch PCR & DNA Cleanup Kit (NEB) according to the manufacturer’s instructions. DNA sequencing libraries were then prepared using the NEBNext Ultra II DNA Library Prep Kit for Illumina according to the manufacturer’s instructions. Each library was purified on 8% TBE gels (Invitrogen), quantified using the KAPA library quantification kit (Roche) and sequenced on an Illumina HiSeq 4000 with 2 × 150 bp paired-end runs.

### Deriving mean stall scores

The reads associated to the stall products were pre-processed with cutadapt [37] to trim the dC-tails using a quality cut-off of 20 and identifying tails with a minimum of 10 consecutive Cs (or Gs on the reverse reads). Reads were mapped to an artificial library chromosome using Bowtie 2 [38] using the local and end-to-end functions for the stalled and extended products respectively. A minimum of 67 and 88% of overall alignment rates were obtained for the stalled and extended products respectively with a minimum of 6.5 million aligned reads per library. We obtained the number of mapped reads to each sequence from the library using the idxstats command of Samtools [39]. When a given sequence has no reads associated to it in a given condition, an arbitrary value of one over the total number of reads obtained for this library was assigned to it. The number of reads for each sequence for each condition was then normalised by dividing its value by the total number of mapped reads at this condition and its abundance expressed in counts per million (CPM). Stall scores were then computed as the ratio of the number of reads associated with a sequence in the stall fraction over the total number of reads associated to the same sequence in both the stalled and extended fractions. Hence, the 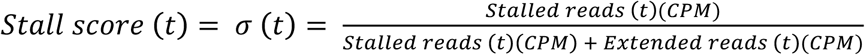 is a number between 0 and 1 with values of 0 describing sequences that are not present in the stalled fractions and values of 1 describing sequences found exclusively in the stalled fractions. Means and standard deviations were computed from values obtained from duplicates, *i*.*e*. using ssDNA templates from two independent M13KO7 helper phage preparations. Sequences with standard deviation of stall scores higher than the 95 percentile, *i*.*e*. outliers were excluded from the analysis.

### Kinetics of resolution of DNA polymerase stalling

As a proxy for the kinetics of resolution of a stall event at each sequence from the library we analysed the variation of stall scores overtime and extract a kinetic constant (λ) using an exponential growth/decay model: σ (*t*) = σ_0_. *e*^*λt*^. This model was chosen due to the known exponential behaviour of the association and dissociation kinetics of single DNA-synthesising T7 DNA polymerase [29]. In this context, positive and negative values for λ reflect persistent or transient stalling events respectively.

### Structure features

To assess the correlation between the stall scores and the stability of DNA structures, we predicted their stability as follows: (i) for hairpin structures we predicted a melting temperature *Tm* based on the formula [40] *Tm* = *L* × (*2*×%*GC* + 2), where *L* is the length of the stem of the hairpin and %GC the GC content of the sequence, (ii) for tetrahelical structures we computed a G4Hscore which is a quantitative estimation of G-richness and G-skewness that correlate with the folding propensity [41]. Briefly, each position in a sequence is given a score between −4 and 4. To account for G-richness, a single G is given a score of 1, in a GG sequence each G is given a score of 2; in a GGG sequence each G is given a score of 3; and in a sequence of 4 or more Gs each G is given a score of 4. To account for G-skewness, Cs are scored similarly but values are negative. While high positive G4Hscore indicate G4 formation, low negative values indicate i-motifs formation. To assess sequence diversity within STRs, we computed the Shannon entropy *H* of each motif according to the formula: *H* = − Σ *f*_*a,i*_ × *log*_2_ *f*_*a,i*_, where *f*_*a,i*_ is the relative frequency of each base a at position i.

### STR structure classification using a supervised machine learning approach

The selection of a classification algorithm and the prediction of STR structures were performed using the “caret” package [42] in the *R* (https://www.R-project.org/) environment. The set of 2,932 control sequences was used to train, test and select the best performing algorithm. Each of the sequences was classified into one of the four structural classes HAIRP, IMOT, QUAD and UNF for hairpins, i-motifs, G4s and unfolded sequences respectively. UNF sequences comprised sequences whose stall scores are not statistically different from the scores of the negative control sequences, *i*.*e*. random sequences of varying GC content, at each time points. To identify those sequences, we assign to each stall scores a *P* value at each time point, using a Mann-Whitney U test challenging the replicate values against the distribution of values obtained for the negative control sequences, and combining these *P* values according to Fisher’s method [43]. Sequences with combined *P* values, referred to as *Q*-values, higher than 0.1 were defined as UNF. The set of sequences used to train the classifiers then comprised 427 HAIRP, 105 IMOT, 983 QUAD and 1,417 UNF sequences. The set of sequences was then randomly portioned into two sets: 70% of the sets were used for training and the remaining 30% were used for testing. To train the classifiers we considered 11 features, which were the values of the stall scores at each five time point and their associated Q-value, the kinetic constants of stalling resolution λ and their associated coefficient of determination *R*^2^, the GC content, G4Hscore and entropy of the sequences. Each feature was centred and scaled. The distributions of each of the features within the different structural classes are reported Additional file 1: Figure S3a. The training set was used to select models using a k-fold cross validation approach (“cv” model from the caret package) with 10 numbers of folds. To assess the overall performance of the models, we then challenged them against the test set. The model performing with higher accuracy, *i*.*e*. a random forest algorithm performing with an accuracy of 0.96 ± 0.03 over 100 resamplings, was selected to predict the structure of STR motifs. It is noteworthy that a similar model selected using only features relative to sequence features, *i*.*e*. GC content, G4Hscore and entropy, is performed with an accuracy of 0.64 ± 0.13 over 100 resamplings (Additional file 1: Figure S3c), indicating that information about polymerase stalling is essential for assigning structures. Each of the 5,356 possible permutations of 1-6 nt long STR sequence was then assigned into 964 unique single-stranded motifs. Average stall scores, *P* values and combined *Q*-values were computed, as previously described, and each of the STR motifs was classified into the four different structural classes for three different lengths. Structural assignation of each STR can be found in Additional file 2.

### Hierarchical clustering of STR motifs

To assess the hierarchical sequence relationship between STR motifs, similarities between motifs were assessed using a cosine similarity score. A cosine similarity score between two sequences A and B is computed as 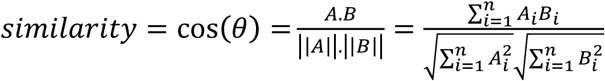, where *A*_*i*_ and *B*_*i*_ are the components of the *N*-gram (N = 2) vectors *A* and *B* associated to both motifs. *N*-gram vectors of length *n*, associated to motifs A and B, were constructed by computing the number of occurrences with consecutive *k*-mers (*k* = 2) present in both motifs. Cosine similarity was chosen due to its sensitivity for mapping noisy and repetitive sequences [44]. A dendrogram based on distances computed from cosine similarities was then generated using the “dendextend” package [45] in the *R* environment. Finally, the branches of the tree were coloured according to the predicted structure of the STR motifs when 72-nt long.

### NMR spectroscopy

The HPLC-purified oligonucleotide sequences used for NMR spectroscopy are shown in Additional File 1: Table S1. 1H NMR spectra were recorded at 298 K using a 600-MHz Bruker Avance I spectrometer equipped with a cryogenic TCI probe. Water suppression was achieved using a WATERGATE pulse-sequence modified in-house with a water flipback element for optimal solvent suppression. Watergate spectra were also collected with or without an additional presaturation pulse when required. The oligonucleotides were annealed at a final concentration of 0.2 mM in the same buffer used for the primer extension assay which is 40 mM Tris.HCl pH 7.5, 20 mM MgCl_2_, 50 mM NaCl and 50 mM KCl supplemented with 10% D_2_O. The samples were annealed by heating at 80 °C for 10 min and slowly cooled to 4 °C.

### Meta-sequence coverage profiles

Coverage data from estimated counts, were generated using the deepTools software [46] and the bamCoverage command with a bin size of 1. To generate normalised profiles of the stalled products, coverage values at each position for each sequence were divided by the maximum values observed for this sequence. The means of the normalised counts at each position from both duplicates were then computed to generate the relative coverage profiles. Meta-sequence coverage profiles were then generated by considering the average values at a given position across the set of sequences of interest. The positions of the stalled polymerase were defined as the position at which the relative coverage is equal at 0.5. In order to assess the distance travelled by the polymerase within the STRs, the difference between the positions of the stalled polymerase at a given time and the position at 0.5 min was considered.

### Sequence variants calling

Sequence variants were called with FreeBayes [47] which is an haplotype-based Bayesian statistical framework for detection of base substitutions and indels. BAM files from duplicates were merge in order to increase the number of reads per STRs and the sensitivity of the analysis. Variants calling was performed on the extended products at each time point and on the parental library, *i*.*e*. the library that hasn’t been extended by the T7 DNA polymerase, using the --ploidy 1 --use-duplicate-reads --no-complex options of FreeBayes. Duplicates reads were used because all sequencing reads are in fact amplicons. Base substitutions (SNP) and expansion/contraction (INDEL) were then considered independently. We applied a series of filter to select only variants of interest. We first selected SNPs and INDELs with estimated phred-scaled base quality above 20. To select *de novo* mutations, *i*.*e*. mutations arising from DNA synthesis by the T7 DNA polymerase rather than artefacts originating from the cloning, PCR and library preparation steps, any mutations called within the parental library were then excluded from the analysis. To define the universe of observable mutations, all the mutations called at each time points and within the repeats were finally combined and considered for further analysis. For expansion/contraction events only INDELs of sizes equal to the multiples of the unit size and of the same sequence were considered. The frequency of base substitutions and expansion/contraction events was computed by dividing the number of reads supporting a mutation by the total number of reads covering this mutation.

### Abundance and length of eukaryotic STRs

Genomic coordinates of STRs from 6 eukaryotic genomes were recovered from the MicroSatellite DataBase [3]. The analysed genomes were *Homo sapiens* (hg38), *Mus musculus* (mm10), *Gallus gallus* (galGal6), *Danio rerio* (dm6), *Drosophila melanogaster* (dm6) and *Saccharomyces cerevisiae* (sacCer3). The MicroSatellite DataBase reports perfect STRs identified with the PERF algorithm. To assess the correlation between stall scores, abundance and length, each 5,356 possible permutations of 1-6 nts long STR sequences were classified into 501 unique double-stranded motifs and the average stall scores were considered for the analysis. Mononucleotide repeats were excluded from the analysis due to their known overrepresentation in eukaryotic genomes [48].

### STR mutation rates and length constraints

Expansion and contraction rates (μ), the strength of the directional bias of mutation (*β*) and the geometric mutation step size distribution (*p*) of individual STRs was recovered from reference [24], reporting mutation rate estimates for 1,250,930 autosomal human STRs. We excluded from the analysis mononucleotide repeats and repeats shorter than 12 nucleotides to ensure at least two repeats of hexanucleotide motifs. To assess any correlation between STR stall scores and mutability parameters, we classified each permutation of STR sequences into 501 unique double-stranded motifs and average stall scores were considered for the analyses. We considered only STR loci displaying mutation rates above 10^−7.5^ events per generation, which is the lower limit of quantification of mutation rates by this model.

### Common sequence variation at STR loci

All common germline variants from dbSNP build 150 on hg19 (common_all_20170710.vcf.gz) were recovered from ftp://ftp.ncbi.nih.gov/snp/organisms/human_9606_b150_GRCh37p13/VCF/. This dataset reports all variants representing alleles observed in the germline with a minor allele frequency ≥ 0.01 in at least one 1000 Genomes Phase III major population, with at least two individuals from different families having the same minor allele, and consists of 34,082,223 SNPs. We selected these well-characterized SNPs in order to ensure that the correlations drawn in our work are not due to sequencing artefacts. SNP frequency at individual positions was computed by dividing the number of STR marked by a SNP at a given position by the total number of considered STRs. SNP enrichment in the vicinity of STR was assessed using two-sided Fisher tests considering the average frequency of SNPs at position −1 and +1 from the start and the end of the repeats. Nucleotide substitution preferences at internal positions of the repeats (from the start to start + 6 nt and from the end – 6 nt to the end) were used to build the substitution matrices associated to each STR structural class.

### Statistics

In relevant figures, figure legends convey the statistical details of experiments, while asterisks define degree of significance as described. All statistical analyses were performed under the *R* environment.

## Additional files

**Additional file 1: Supplementary Information comprising: Figure S1**. Primer extension assay quality controls. **Figure S2**. The kinetics of DNA synthesis highlights structure-dependent transient and persistent stalling events. **Figure S3**. Supervised machine learning approach to structure prediction from DNA polymerase stalling events. **Figure S4**. Inferring STR structures from DNA polymerase stalling events. **Figure S5**. Validation of predicted STR structures. **Figure S6**. The DNA polymerase remodels STRs during DNA synthesis. **Figure S7**. Identification of sequence variants within the pool of newly synthesised DNA molecules. **Figure S8**. STR structures impact the frequency and nature of expansion/contraction events. **Figure S9**. STR structures impact the frequency and nature of nucleotide substitution events. **Figure S10**. Structure-dependent nucleotide substitution preferences. **Figure S11**. DNA polymerase stalling at DNA structures predicts abundance and length of STRs in the human genome. **Figure S12**. DNA polymerase stalling at DNA structures predicts abundance and length of STRs in eukaryotic genomes. **Figure S13**. DNA polymerase stalling at DNA structures predicts STR stability in the human genome. **Figure S14**. Structured STRs are prone to sequence variation in the human genome. **Table S1**. Oligonucleotides used for the validation of STR structures by ^1^H NMR spectroscopy.

**Additional file 2:** Sequences, stall scores, kinetics parameters and predicted structures of the 20,000 designed DNA sequences (CSV file).

## Availability of data and materials

The data that support the findings of this study are available from Additional file 2 and from the corresponding authors upon request. Sequencing data can be accessed at the Gene Expression Omnibus archive with the accession number GSE144458. The codes and materials used in this study are available from the corresponding authors upon request.

## Author contributions

PM designed the project and performed the experiments. All authors analysed and interpreted the data. PM and JES wrote the manuscript.

## Acknowledgements

The authors thank J. Wagstaff and J.-C. Yang for help with NMR spectroscopy experiments and M. Babu, J. Yeeles and the members of the Sale group for comments on the manuscript.

## Ethics approval and consent to participate

Not applicable.

